# Evaluation of a Novel Recombinant Human Protein Formula Compared to Donor Human Milk and Standard Formula in Neonatal Piglets

**DOI:** 10.64898/2026.08.12.744500

**Authors:** Sarah Elefson, Valeria Melendez Hebib, Gregory Hoeprich, Jonathan Lau, Julia de Macedo Robert, Mariana Wanessa Santana de Souza, Mauro Ramalho Silva, Caitlin Vonderohe, Gregory Guthrie, Barbara Stoll, Del Alfonso, Douglas Burrin

## Abstract

**Background:** Despite the advancements in infant nutrition, a gap still exists in the nutritional composition bioactive ingredients between infant formula and human milk. We developed a next-generation, proof-of-concept infant formula that contains recombinant human milk proteins.

**Objective:** To determine the impact of a novel infant formula (H1) on organ growth and development, and intestinal function compared to donor human milk (DHM) and standard infant formula (S) in a term piglet model.

**Methods:** Term piglets delivered via cesarean section were fed either a donor human milk (DHM) control, the investigational formula (H1), or infant formula (S) for 10 days. On d 10, a blood sample and tissues were collected.

**Results:** There was no difference (*P* > 0.05) in piglet growth, although H1 piglets had a smaller relative stomach and liver than DHM and S piglets. H1 piglets had higher (*P* < 0.05) interleukins in the distal ileum, but no other systemic cytokines were elevated compared to the DHM and S piglets. H1 piglet small intestinal histology was similar (*P* > 0.05) to that of DHM and S piglets. Additionally, H1 piglets had either the same (*P* > 0.05) or higher (*P* < 0.05) amino acids in circulation compared to DHM and S piglets. Recombinant human proteins had either similar (*P* > 0.05) or lower (*P* < 0.05) activity compared to the native human proteins when assessing the individual ingredients in the H1 formula.

**Conclusion:** H1 formula was noninferior to DHM and S based on growth, small intestinal histology and plasma amino acid endpoints when fed to neonatal piglets. These findings warrant further studies to use the neonatal piglet as a model to evaluate more in-depth outcomes of health and safety for new infant formulas.

**Lay Summary:** A novel piglet study shows a hypoallergenic, next-generation infant formula containing recombinant human milk proteins rivals donor human milk and standard formula for growth, gut health, and nutrient status.

## Introduction

Human milk is considered the gold standard recommended diet for infants in the first six months of life, yet only ∼55% receive breast milk and thus, substantial numbers of infants receive infant formula (1, 2). Human milk contains a complex combination of macro- and micronutrients, hormones, cytokines, and human milk oligosaccharides (HMOs) (3). Infant formulas that are used to complement or substitute for human milk are made primarily of cow’s milk, other animal, and/or vegetable products (4). Several ingredients in human and cow’s milk, especially proteins, not only have nutritional value to provide amino acids but also have functional properties involved in digestion and immune function (4, 5).

The prevalence of food allergies has increased over the past few decades. Allergies often arise in infants and children under three years of age, with incidence rates reported around 8% of individuals being affected (6, 7). Infants who develop a food allergy are restricted in which formulas they can use safely, and it can even limit the diet of a woman who is breastfeeding, as proteins will transfer to their milk. Thus, the elimination of bovine and soy proteins in formula can help to reduce allergic reactions that could occur in infants.

There is a need to close the gap in nutrient composition and function between infant formulas and human milk (8). In this study, we evaluated a new formula developed with recombinant human proteins, including albumin, lactoferrin and lysozyme, plus free amino acids to provide protein, human milk oligosaccharides (HMO) and all other nutrients required by infants. This next-generation proof-of-concept infant formula allows for a more similar breastmilk composition and is hypoallergenic. The objective of this study was to determine the impact of a novel formula on organ growth and development, and intestinal integrity compared to donor human milk (DHM) and infant formula in a piglet model. Development of new pre-clinical models to evaluate the safety and nutrient bioavailability of new infant formulas is important (9). The neonatal piglet is a well-developed translational model for studying nutrition and metabolism as would occur in human infants (10-14). Due to the similarity between neonatal piglets and human infants, particularly as it relates to their physiology and metabolism, we consider the piglet to be the most appropriate model to investigate the safety and nutritional effects of infant formula components. We included a treatment group fed donor human milk as reference control in addition to a comparison group fed a commercial infant formula. Additionally, components of the novel infant formula were measured for activity to determine comparability to human proteins. We hypothesized that H1 is noninferior when compared to donor human milk and commercial infant formula.

## Materials and Methods

### Animal Care and Feeding

The animal protocol was approved by the Institutional Animal Care and Use Committee of Baylor Medicine and was conducted in accordance with the National Institute of Health guidelines. One pregnant sow (commercial landrace mix breed) was transported to the facility one week prior to the start of the study for acclimation. Piglets (N=18) were delivered via cesarean section one day before the estimated term due date (gestation day 114/115).

Piglets were surgically implanted with a jugular venous catheter (0.03″ ID, silastic) and an orogastric tube (6 French Tygon) to facilitate parenteral nutrition and enteral feeding as previously described (15, 16). Following surgery, these pigs received 16 mL/kg maternal plasma intravenously in 3 doses within 24 hours to provide passive immunity. Pigs were individually housed in 2 ft^2^ stainless steel cages with tenderfoot flooring for three weeks under a 12-hour light-dark cycle at 31-32°C. Pigs received 50% of their full nutritional requirements (240 kcal, 13 g amino acid, 25 g glucose, 5 g lipid, and 240 mL fluid) via total parental nutrition at a rate of 5 mL fluid/kg per hour d 1, 2, and 3. Starting d 2, enteral feeds were administered at 25% of their nutrition requirement. The volume of enteral feeds increased to 100% by d 5, and 125% by d 10 (**Figure 1**), to maximize nutrient intake to meet the higher requirements of neonatal pigs (**Table 1)**. Pigs were randomized to receive either donor human milk (DHM) that was pasteurized using the Holder pasteurization method, term infant formula (S) (Similac; Abbott, Chicago, IL), or the test novel infant formula (H1) (**Table 2**). At the end of the 10-day study, piglets were humanely euthanized.

**Figure 1.**
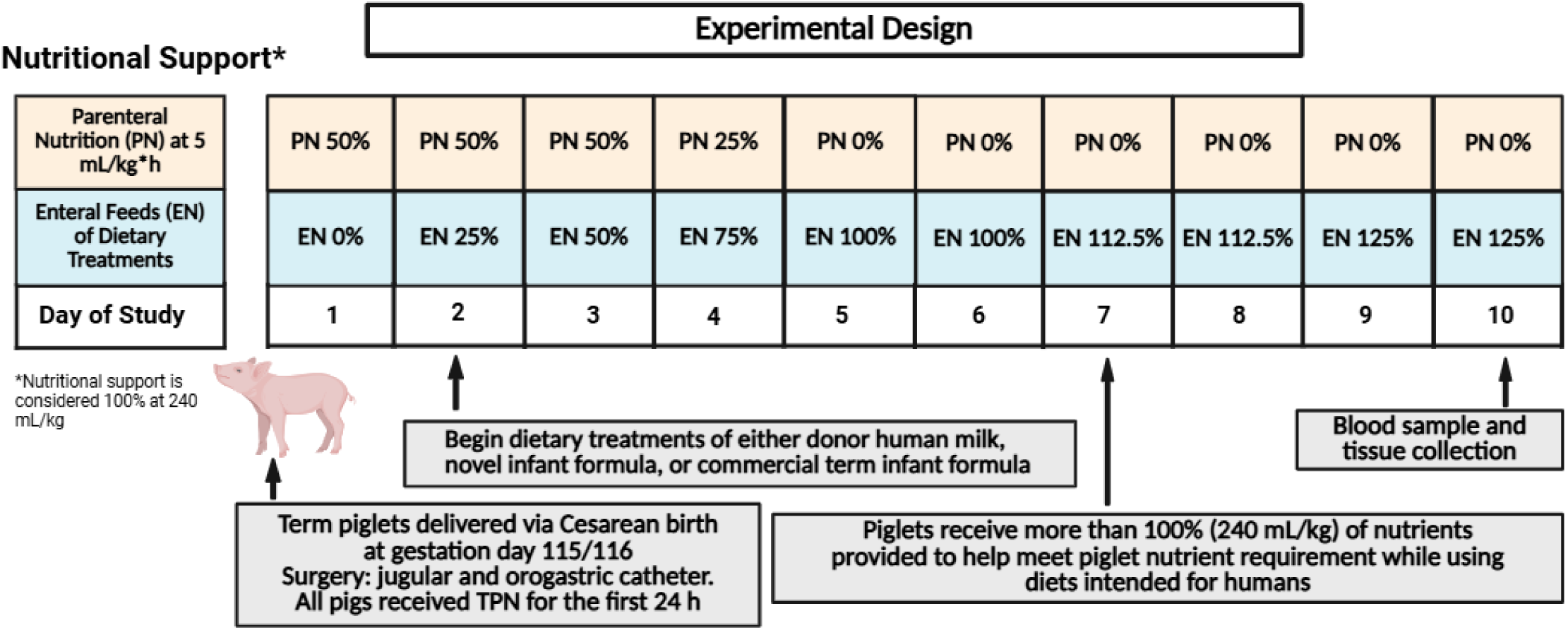
Experimental Study Design. Created with BioRender.

**Table 1.**
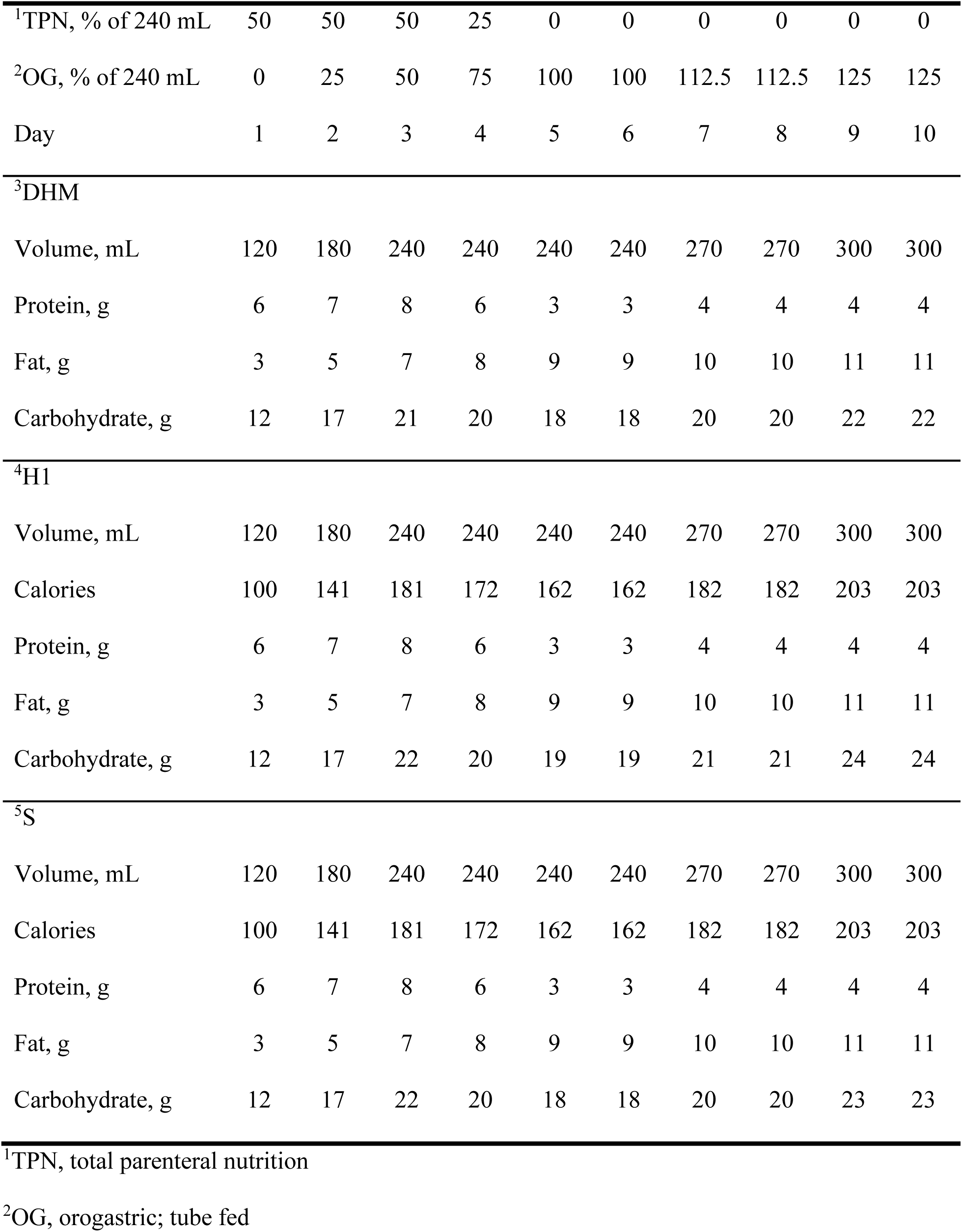
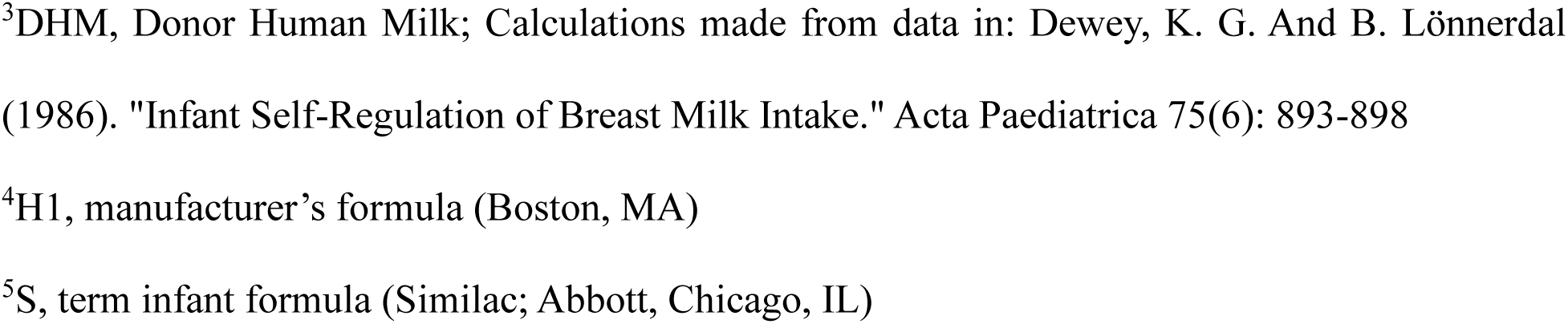
Calculates nutrient content of dietary treatments.

|  |  |  |  |  |  |  |  |  |  |  |
| --- | --- | --- | --- | --- | --- | --- | --- | --- | --- | --- |
| <sup>1</sup>TPN, % of 240 mL | 50 | 50 | 50 | 25 | 0 | 0 | 0 | 0 | 0 | 0 |
| <sup>2</sup>OG, % of 240 mL | 0 | 25 | 50 | 75 | 100 | 100 | 112.5 | 112.5 | 125 | 125 |
| Day | 1 | 2 | 3 | 4 | 5 | 6 | 7 | 8 | 9 | 10 |
| <sup>3</sup>DHM |  |  |  |  |  |  |  |  |  |  |
| Volume, mL | 120 | 180 | 240 | 240 | 240 | 240 | 270 | 270 | 300 | 300 |
| Protein, g | 6 | 7 | 8 | 6 | 3 | 3 | 4 | 4 | 4 | 4 |
| Fat, g | 3 | 5 | 7 | 8 | 9 | 9 | 10 | 10 | 11 | 11 |
| Carbohydrate, g | 12 | 17 | 21 | 20 | 18 | 18 | 20 | 20 | 22 | 22 |
| <sup>4</sup>H1 |  |  |  |  |  |  |  |  |  |  |
| Volume, mL | 120 | 180 | 240 | 240 | 240 | 240 | 270 | 270 | 300 | 300 |
| Calories | 100 | 141 | 181 | 172 | 162 | 162 | 182 | 182 | 203 | 203 |
| Protein, g | 6 | 7 | 8 | 6 | 3 | 3 | 4 | 4 | 4 | 4 |
| Fat, g | 3 | 5 | 7 | 8 | 9 | 9 | 10 | 10 | 11 | 11 |
| Carbohydrate, g | 12 | 17 | 22 | 20 | 19 | 19 | 21 | 21 | 24 | 24 |
| <sup>5</sup>S |  |  |  |  |  |  |  |  |  |  |
| Volume, mL | 120 | 180 | 240 | 240 | 240 | 240 | 270 | 270 | 300 | 300 |
| Calories | 100 | 141 | 181 | 172 | 162 | 162 | 182 | 182 | 203 | 203 |
| Protein, g | 6 | 7 | 8 | 6 | 3 | 3 | 4 | 4 | 4 | 4 |
| Fat, g | 3 | 5 | 7 | 8 | 9 | 9 | 10 | 10 | 11 | 11 |
| Carbohydrate, g | 12 | 17 | 22 | 20 | 18 | 18 | 20 | 20 | 23 | 23 |

<sup>1</sup>TPN, total parenteral nutrition
<sup>2</sup>OG, orogastric; tube fed
<sup>3</sup>DHM, Donor Human Milk; Calculations made from data in: Dewey, K. G. And B. Lönnerdal (1986). "Infant Self-Regulation of Breast Milk Intake." *Acta Paediatrica* 75(6): 893-898 <sup>4</sup>H1, manufacturer's formula (Boston, MA) <sup>5</sup>S, term infant formula (Similac; Abbott, Chicago, IL)

**Table 2.**
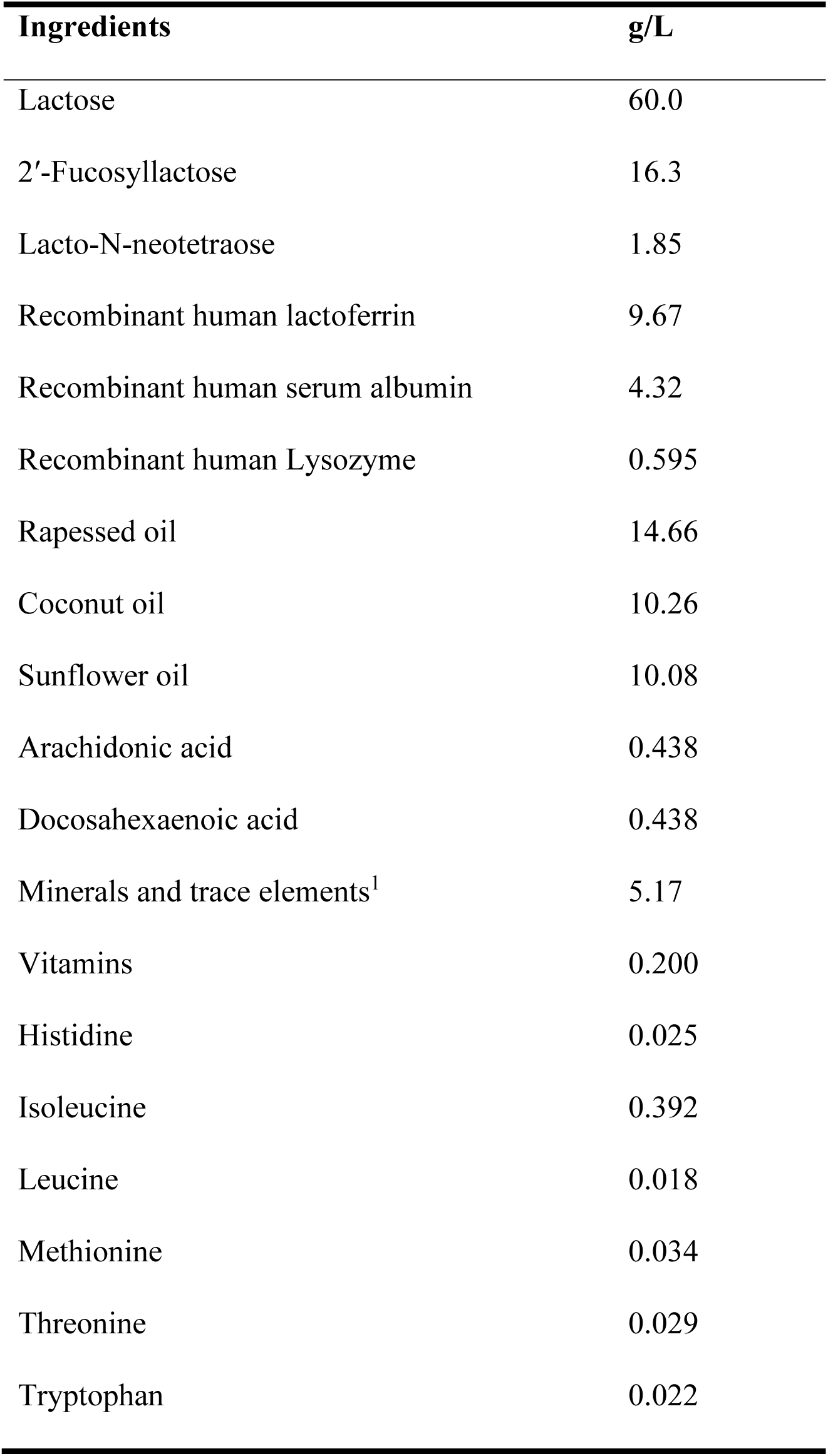
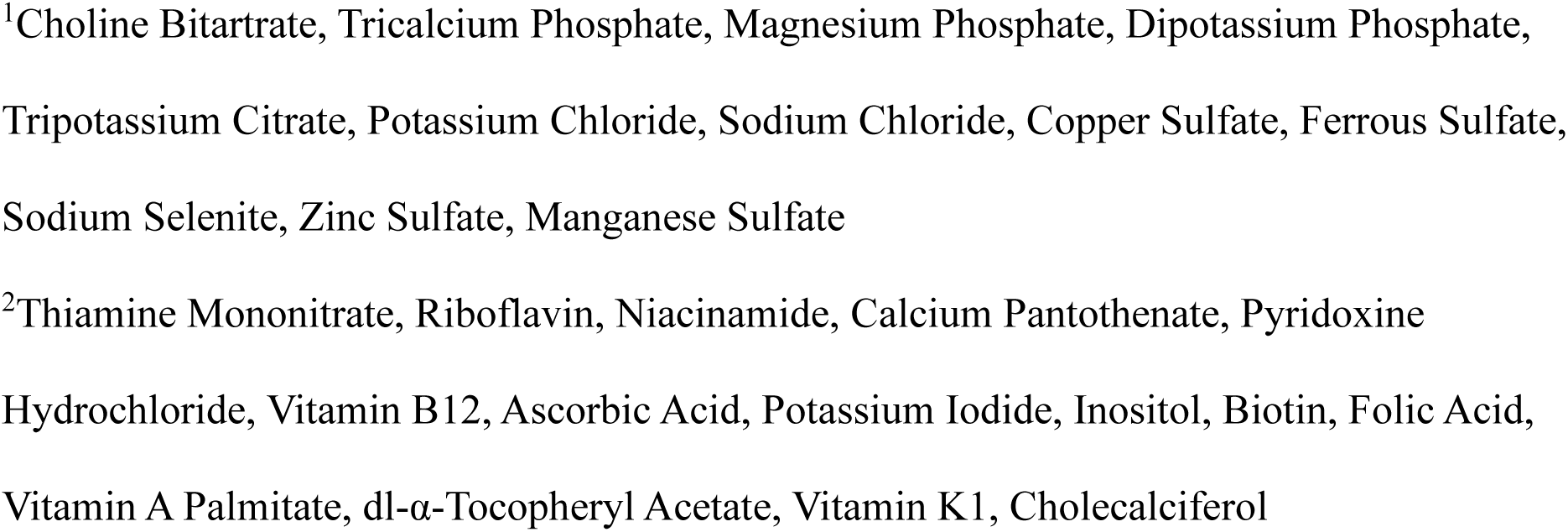
H1 Ingredient Composition.

| **Ingredients** | **g/L** |
| --- | --- |
| Lactose | 60.0 |
| 2'-Fucosyllactose | 16.3 |
| Lacto-N-neotetraose | 1.85 |
| Recombinant human lactoferrin | 9.67 |
| Recombinant human serum albumin | 4.32 |
| Recombinant human Lysozyme | 0.595 |
| Rapessed oil | 14.66 |
| Coconut oil | 10.26 |
| Sunflower oil | 10.08 |
| Arachidonic acid | 0.438 |
| Docosahexaenoic acid | 0.438 |
| Minerals and trace elements<sup>1</sup> | 5.17 |
| Vitamins | 0.200 |
| Histidine | 0.025 |
| Isoleucine | 0.392 |
| Leucine | 0.018 |
| Methionine | 0.034 |
| Threonine | 0.029 |
| Tryptophan | 0.022 |

<sup>1</sup>Choline Bitartrate, Tricalcium Phosphate, Magnesium Phosphate, Dipotassium Phosphate, Tripotassium Citrate, Potassium Chloride, Sodium Chloride, Copper Sulfate, Ferrous Sulfate, Sodium Selenite, Zinc Sulfate, Manganese Sulfate <sup>2</sup>Thiamine Mononitrate, Riboflavin, Niacinamide, Calcium Pantothenate, Pyridoxine Hydrochloride, Vitamin B12, Ascorbic Acid, Potassium Iodide, Inositol, Biotin, Folic Acid, Vitamin A Palmitate, dl- $\alpha$ -Tocopheryl Acetate, Vitamin K1, Cholecalciferol

### Sample Collection

Body weight and plasma samples were taken every other day beginning on d 1 of the study. Blood samples were separated for either serum or plasma, via centrifugation at 1,500 g for 10 minutes at 4°C, then frozen at -80°C until further analysis. After the piglets were euthanized on d 10, tissues were dissected, weighed, and sections of selected organs were snap-frozen or fixed in either formalin or Optimal Cutting Temperature compound for later analysis. All biological samples were kept at -80°C until analysis was conducted.

### Plasma Analysis

Plasma samples were measured for complete blood count (CBC), including white blood cells (WBC), neutrophils, lymphocytes, and eosinophils through a core facility at Baylor College of Medicine.

### Cytokine and Amino Acid Analysis

Plasma, liver, and distal ileum were analyzed using the porcine Cytokine/Chemokine Discovery Assay using Luminex platform by EVE technologies (Calgary, Canda). Samples were prepared according to EVE’s protocols and normalized to 3 ng/mL protein per aliquot. Amino acid analysis of plasma was conducted using liquid chromatography-mass spectrometry.

### Immunohistology Small Intestine Histology Analysis

Subsections of the liver were stained for Ki67 to measure proliferation of cells, while subsections of the distal ileum were stained with alcian blue to detect mucus in the small intestine as described previously (17). The histology slides were scanned and then analyzed with Image J (18) by separating color channels to then analyzing the percent of appropriate color present on the section of the slide via set thresholds. Thresholds were set to optimally capture the staining on the slides, and then the average percent of the stain was recorded and analyzed. Sections of the proximal jejunum and distal ileum were stained with hematoxylin and eosin to determine inflammation. The inflammation score was completed by a board-certified veterinarian who was blinded to treatments. Scores were recorded for inflammation in the lamina propria, number of eosinophils present, percent of goblet cells, and vacuole presence.

### H1 Recombinant Human Protein Analysis

#### Esterase Activity (Human Serum Albumin)

The esterase activity of human serum albumin (HSA) was determined by using a colorimetric assay. This involved the cleavage of an ester bond in the enzyme substrate to generate a yellow product. A stock solution of 30 mM solution of p-nitrophenyl acetate (Sigma N8130) was prepared in DMSO:methanol (1:1 v/v). It was diluted to a 3 mM working stock with 20 mM Tris-buffered saline (TBS) at the time of the assay. For a standard curve to measure how many moles of product was formed, a stock solution of 30 mM p-nitrophenol was prepared in DMSO:methanol (1:1 v/v). It was diluted to a 3 mM working stock with 20 mM TBS, followed by serial dilution to achieve concentrations in the range of 0 – 100 uM. The samples compared were: Oryzogen recombinant HSA (HYC002R02); Roche’s bovine serum albumin fraction V (0735078001); and Sigma HSA standard (A3782). The albumin proteins were reconstituted at a concentration of 1 mg/mL in 20 mM TBS. In order to determine the activity of the protein, 25 μL of 3 mM p-nitrophenyl acetate was added to all reaction wells. For normalization, 100 μL of 20 mM TBS was added to indicated wells. For albumin esterase activity, 100 μL of albumin sample was added to indicated wells. The 96 well plate was entered into the plate reader and the protocol was started. This protocol involved a brief shake, followed by reads at 1, 2, 3, and 6 min at 400 nm to determine the absorbance. To account for the spontaneous degradation of p-nitrophenyl acetate, all albumin containing wells were normalized with the control well (TBS only). Using the p-nitrophenol standard curve, the following equation was then used to calculate the esterase activity:

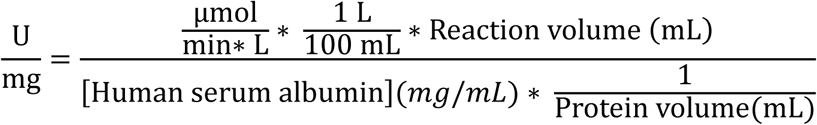

### Lysozymes Activity Assay

Lysozyme activity was quantified using the EnzChek® Lysozyme Assay Kit (E-22013, Molecular Probes), following the manufacturer’s protocol with minor modifications. The assay is based on the fluorescence increase resulting from the enzymatic cleavage of fluorescein-labeled *Micrococcus lysodeikticus* cell wall substrate, which becomes de-quenched upon lysozyme digestion. The native human lysozyme protein was purchased from Abcam (ab91125) and the recombinant human lysozyme protein was provided by Oryzogen (HYC042R01). The DQ™ lysozyme substrate (Component A) was reconstituted in deionized water to a final concentration of 1 mg/mL and stored at 4 °C, protected from light. A 1000 U/mL lysozyme standard solution was prepared by reconstituting lyophilized lysozyme (Component C) in deionized water. The 1X reaction buffer (Component B; 0.1 M sodium phosphate, 0.1 M NaCl, pH 7.5, containing 2 mM sodium azide) was used throughout the assay. All reactions were conducted in 96-well black-walled microplates in a final volume of 100 μL. Lysozyme standards (ranging from 0 to 250 U/mL final concentration) were prepared by serial twofold dilutions of the 1000 U/mL stock solution in 1X reaction buffer. Experimental samples were similarly diluted in 1X reaction buffer to a total of 50 μL per well.

The working substrate solution was prepared at 50 μg/mL by diluting the 1 mg/mL DQ lysozyme stock 20-fold in 1X reaction buffer. To initiate the reaction, 50 μL of the working substrate solution was added to each well containing 50 μL of sample or standard. Plates were incubated at 37 °C for 30 minutes, protected from light. Fluorescence was measured using a GloMax Discover Plate Reader (Ex: 475 nm, Em: 500-550 nm). Background fluorescence was subtracted using the no-enzyme control. Sample fluorescence values were interpolated from the standard curve generated using known concentrations of lysozyme. All samples were measured in triplicate.

### Iron II Binding on Isolated Protein (HSA)

The quantification of iron II binding by HSA was based on Narmuratova’s (19) assay working with lactoferrin binding of iron II. This method relies upon the reaction of the chromatogen with free (unchelated) iron II in the solution. A 2 mM solution of iron (II) chloride was prepared. A 1 mM solution of ferrozine was prepared as the chromatogen. 0.02 M EDTA was used as a positive control. Albumin samples consisted of native HSA (Sigma A3782), recombinant HSA (HYC002R02), and bovine serum albumin (Roche 10735078001) at a concentration of 2 mg/mL in MilliQ water. These samples were serially diluted to achieve concentrations ranging from 0.0625 – 2 mg/mL. For the positive control, 90 μL of MilliQ water was added to a tube, followed by 10 μL of 0.02 M EDTA. For negative control, 200 μL of MilliQ water was added to the tube. A reaction control involved the addition of 100 μL of MilliQ water. For all albumin samples, 100 μL was added to a tube. A volume of 4 μL of 2 mM iron (II) chloride solution was added to each tube. The tubes were shaken vigorously for 15 s, followed by 5 min incubation time at room temperature. Then, 100 μL of 1 mM ferrozine was added to all tubes except for the negative control. The tubes were shaken vigorously for 15 s, followed by 5 min incubation. All samples were centrifuged at 20,000 x g for 2 min. A total of 100 μL from the aqueous phase was transferred to a well on a 96 well plate, followed by an absorbance measurement at 562 nm. The absorbance values were normalized by subtracting the absorbance of the negative control. Percent chelation was calculated using the following equation:

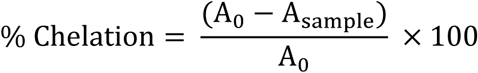

A_0_ is the normalized absorbance for the well with 0 albumin added.

A_sample_ is the normalized absorbance for wells containing sample.

The % chelation values were then plotted against protein concentration and used to determine the protein concentration required to chelate 50% of iron (II).

### Iron III Binding on Isolated Protein (HSA)

The quantification of iron III binding by HSA relies on the reaction of the chromagen with free (unchelated) iron III in the solution. A 20 mM solution of iron (III) chloride tetrahydrate was prepared. A 20 mM solution of potassium thiocyanate was prepared. 0.02M EDTA was used as a positive control. Albumin samples consisted of native human serum albumin (Sigma A3782), recombinant human serum albumin (HYC002R02), and bovine serum albumin (Roche 10735078001) at a concentration of 12 mg/mL in MilliQ water. These samples were then serially diluted to achieve concentrations ranging from 0.25 – 12 mg/mL. A volume of 10 μL of the 20 mM iron (III) chloride tetrahydrate was added to a well. This was followed by the addition of 100 μL of an albumin sample. The well was gently agitated to mix the solutions, then incubated at room temperature for 3-5 min. 100 μL of 20 mM potassium thiocyanate was added. The plate was briefly agitated, then incubated at room temperature for 2 minutes. The absorbance at 480 (chromatogen detection) and 650 nm (reference) was measured.

After normalizing A480 by subtracting A650, the values were plugged into the following equation:

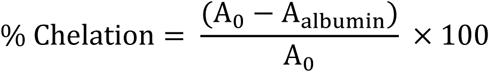

A_0_ is the normalized absorbance for the well with 0 albumin added.

A_albumin_ is the normalized absorbance for wells containing albumin.

The % chelation values were plotted against protein concentration and used to calculate the protein concentration required to chelate 50% of iron (III).

### Lactoferrin Iron II Chelation

The quantification of iron II binding by lactoferrin was based on Narmuratova’s assay (19) working with lactoferrin binding of iron II. This method relies upon the reaction of the chromatogen with free (unchelated) iron II in the solution. A 2 mM solution of iron (II) chloride was prepared. A 1 mM solution of ferrozine was prepared as the chromatogen. 0.02 M EDTA was used as a positive control. Protein samples consisted of Bovine lactoferrin from Glanbia (20: 1058144), bovine lactoferrin from Ingredia (20: SE40277), human recombinant lactoferrin from Oryzogen (20: HYC004M01), Sigma-Aldrich bovine lactoferrin from colostrum (Colostrum, L4765), and Sigma-Aldrich bovine lactoferrin from milk (Milk, L9507). To increase the absorbance range and sensitivity of the assay, the protein samples were reconstituted at a concentration of 2 mg/mL in MilliQ water. These samples were serially diluted to achieve concentrations ranging from 0.0625 – 1 mg/mL. For the positive control, 90 μL of MilliQ water was added to a tube, followed by 10 μL of 0.02 M EDTA. For negative control, 200 μL of MilliQ water was added to the tube. A reaction control involved the addition of 100 μL of MilliQ water. For all albumin samples, 100 μL was added to a tube. A total of 4 μL of 2 mM iron (II) chloride solution was added to each tube. The tubes were shaken vigorously for 15 s, followed by 5 min incubation time at room temperature. 100 μL of 1 mM ferrozine was added to all tubes except for the negative control. The tubes were shaken vigorously for 15 s, followed by 5 min incubation. All samples were centrifuged at 20,000 x g for 2 min. A volume of 100 μL from the aqueous phase was transferred to a well on a 96 well plate, followed by an absorbance measurement at 562 nm. The absorbance values were normalized by subtracting the absorbance of the negative control. Percent chelation was calculated using the following equation:

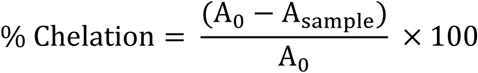

The % chelation values were then plotted against protein concentration and used to determine the protein concentration required to chelate 50% of iron (II) chloride.

### Lactoferrin LPS Binding

A direct enzyme-linked immunosorbent assay (ELISA) was used to assess the binding interaction between purified lactoferrin (hLF) and lipopolysaccharide (LPS) from *Escherichia coli* O111:B4. Plate Coating and Blocking High-binding 96-well ELISA plates (PolySorp, Thermo Scientific, Cat# 456529) were coated with 100 μL per well of LPS (5 μg/mL in 1X phosphate-buffered saline [PBS], Thermo Scientific LPS25) and incubated overnight at room temperature. Plates were washed three times with 200 μL per well of 1X PBST (PBS containing 0.05% Tween-20, Thermo Scientific 28352), followed by blocking with 100 μL per well of 3.3% bovine serum albumin (BSA) in PBS (Thermo Scientific 37525) for 2 hours at room temperature with agitation. Plates were again washed three times with 1X PBST. Purified native human lactoferrin (Abcam, ab285896) or purified bovine lactoferrin (Sigma-Aldrich L9507) was prepared in a twofold serial dilution (ranging from 3.7 μg/mL to 0 μg/mL) in PBST. A volume of 100 μL per well of each concentration was added, and plates were incubated overnight at 4°C. Plates were washed three times with 1X PBST. Primary detection was carried out using a rabbit monoclonal anti-lactoferrin antibody (Abcam, ab259454) for human lactoferrin and HRP anti-lactoferrin antibody (Abcam ab112970) diluted to 1 μg/mL in 1X PBST (100 μL per well). Plates were incubated for 2 hours at room temperature with agitation and washed three times with 1X PBST. Secondary detection was performed with HRP-conjugated goat anti-rabbit IgG (Abcam, ab6721) at a 1:50,000 dilution in PBST (100 μL per well) for human only, followed by 1 hour of incubation at room temperature with agitation. Plates were then washed three times with 1X PBST. A TMB substrate solution (Thermo Scientific, Cat# 34028), pre-equilibrated to room temperature, was added at 100 μL per well. Color development proceeded for approximately 15 minutes at room temperature and was stopped by the addition of 50 μL of 2 M sulfuric acid. Absorbance was measured at 450 nm. All samples were measured in triplicate.

### Statistical Analysis

Data are presented as mean ± standard error of the mean. All data were analyzed in GraphPad Prism v. 10. Except for the small intestinal histology scores, all parameters were analyzed via two-way ANOVA with Tukey adjustments. The heat graphs display the mean of the treatment for each parameter of interest. *P*-values that were *P* ≤ 0.05 were considered significant, while a *P*-value of 0.10 ≥ 0.05 were considered a trend. All graphs were made in GraphPad Prism v 10.

## Results

### Body and Relative Organ Weight

There were no differences in body weight or body weight gain at the end of the study (**Figure 2A-I**). However, piglets that received H1 formula had smaller (*P* < 0.05) relative liver and stomach weights compared to piglets that received DHM and S. There were no other differences in relative organ weights. This indicated that the nutrients provided in the H1 formula are sufficient to support growth in a similar manner to DHM and S (**Table 1**).

**Figure 2.**
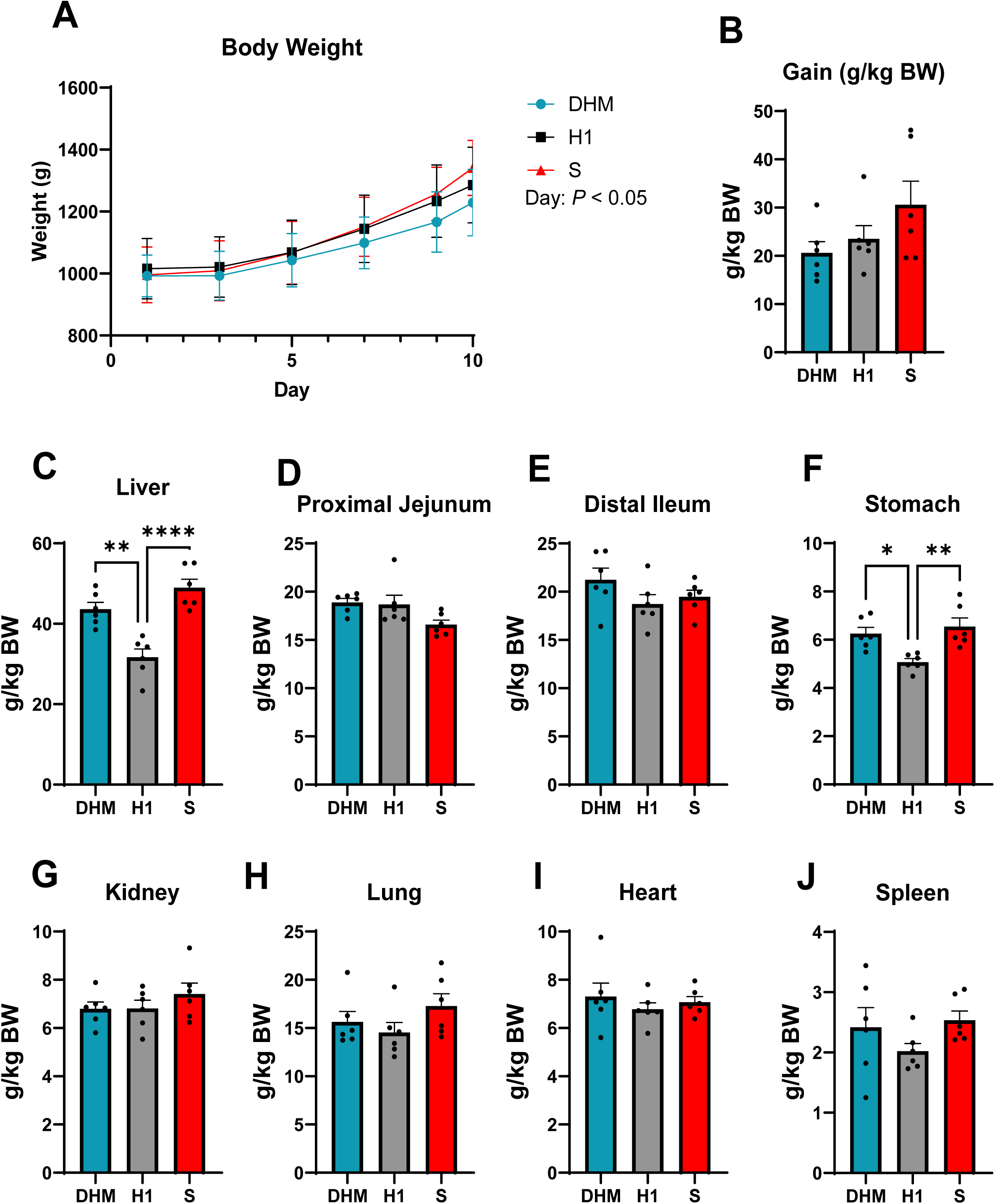
Relative weight gain and organ weights to body weight of piglets on d 10. A-B) Piglet weight gain relative to body weight over ten days with piglets fed either donor human milk (DHM), novel infant formula (H1; Boston, MA, US), or term infant formula (S; Similac; Abbott, Chicago, IL). C-J) Relative organ weight on d 10 of piglets fed either DHM, H1, or S. *, *P* < 0.05; **, *P* < 0.01; ***, *P* < 0.001

### Plasma Complete Blood Count

There were no differences (*P* > 0.05) among the treatment groups for white blood count, neutrophil count, lymphocyte count, and eosinophil count (**Figure 3A-D)**. However, when compared to a piglet reference (21) that establishes intervals for hematology, the piglets who received S diets, had higher (*P* < 0.05) WBC and neutrophils (**Figure 3A** and **Figure 3B**, respectively). These data suggest that H1 does not promote higher inflammation than other dietary treatments. The higher WBC and neutrophils found in piglets that consumed S, could suggest some increased systemic inflammation compared to previously reported values (21).

**Figure 3.**
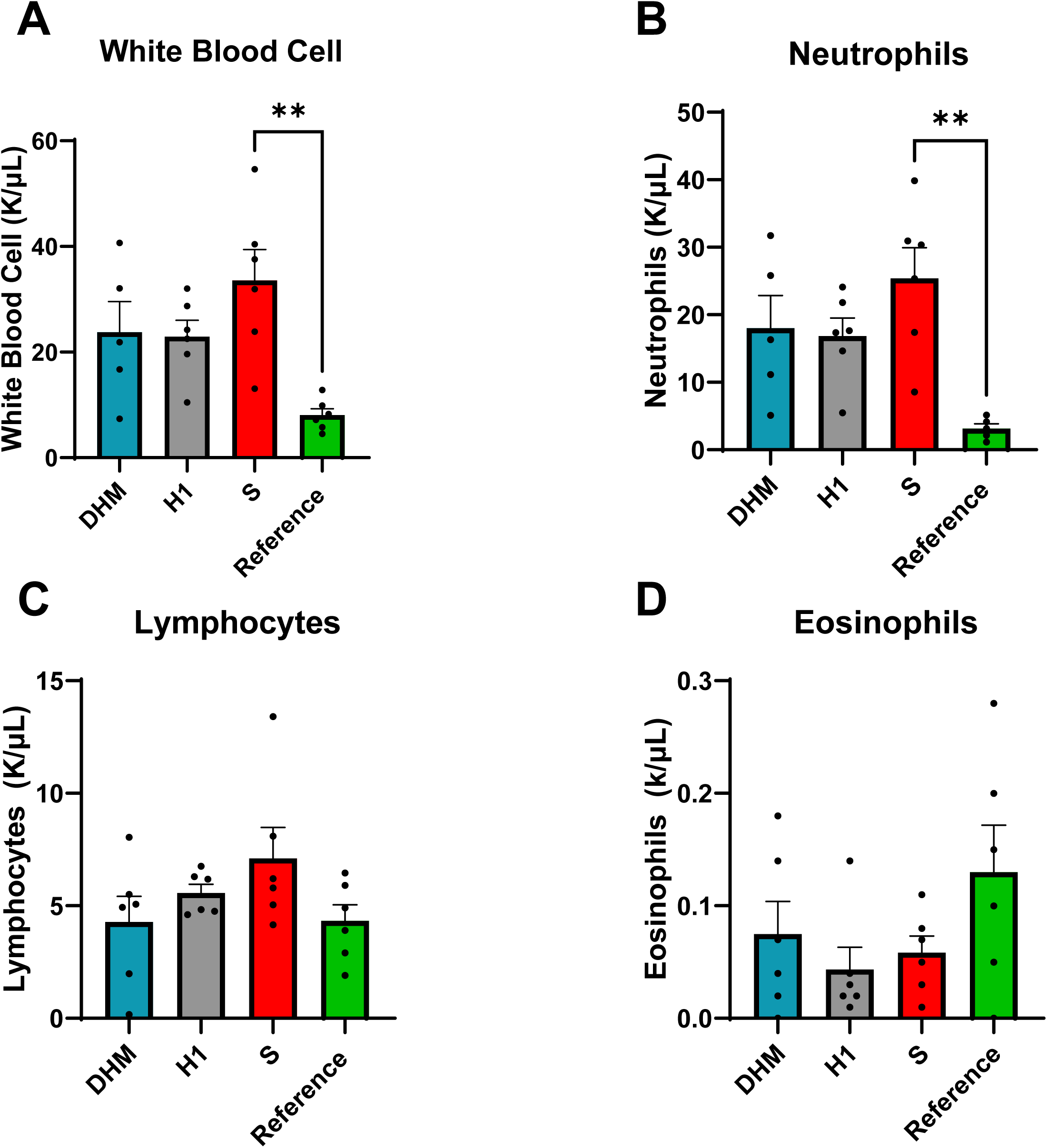
Complete blood cell count of piglets on d 10. A-D) Amount of immune cells present in piglets fed either donor human milk (DHM), novel infant formula (H1; Boston, MA, US), or term infant formula (S; Similac; Abbott, Chicago, IL) on d 10. Reference values from piglets of similar age (21) shown in green bars. *, *P* < 0.05; **, *P* < 0.01; ***, *P* < 0.001

### Small Intestine Histology

Mucin content in the distal ileum as determined by Alcain Blue staining (**Figure 4A**), showed that there were no differences across treatments. DHM and H1 piglets had approximately 95% detection of Alcian Blue while S piglets only had 89.5% detection, although this small difference does not appear to be relevant.

**Figure 4.**
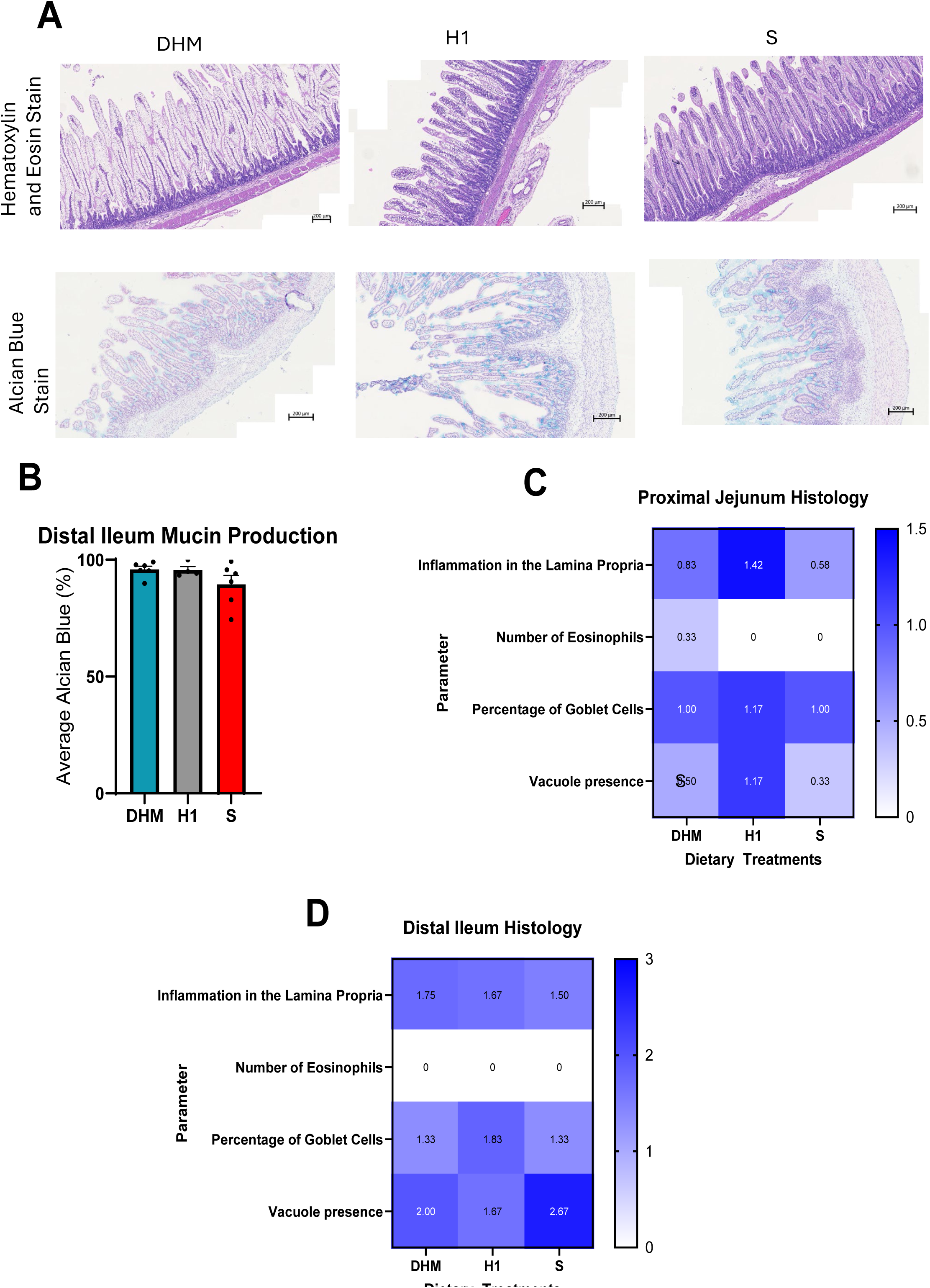
Small intestinal histology staining and small intestinal histology scores of piglets on d 10. A) Representative images of Hematoxylin and Eosin staining from each treatment on d 10. Scale bar, 200 µm. B) Distal ileum mucin production as measured by Alcian Blue staining of term piglets fed either donor human milk (DHM), novel infant formula (H1; Boston, MA, US), or term infant formula (S; Similac; Abbott, Chicago, IL) on d 10. C) Representative images of Alcian Blue staining from each treatment on d 10. Scale bar, 200 µm. F-G) Small intestinal histology scores on d 10 as determined by a board-certified veterinarian.

Histological assessment of the proximal jejunum (**Figure 4C)** showed that lamina propria inflammation scored the highest for H1 pigs (1.42) and lowest for S fed piglets (0.58). Only piglets fed DHM had a detectable score for the number of eosinophils (0.33). There were no statistical differences in mucin content based on goblet cells present, with a mean score of 1.17, while DHM and S pigs had a mean score of 1.00. Piglets receiving H1 had the highest mean score of vacuole presence (1.17), while DHM and S received a lower mean score (0.50 and 0.33, respectively).

In the distal ileum, inflammation in the lamina propria was similar across groups (**Figure 4D)**, with a range of scores from 1.50 to 1.75. All pigs had zero eosinophils present. H1 fed piglets had the highest percentage of goblet cells present, with a mean score of 1.83, while DHM and S pigs had a mean score of 1.33. Pigs receiving S had the highest mean score of vacuole presence (2.67) compared to DHM and H1 piglets (2.00 and 1.67, respectively).

Liver proliferation was measured via a Ki67 stain (**Figure 5A-B**). There were no differences in the amount of Ki67 stained cells between the DHM and H1 piglets; however, H1 piglets had a significantly (P < 0.05) higher amount of Ki67 stained cells compared to S piglets. This result was unexpected considering that H1 piglets had a lower relative liver weight.

**Figure 5.**
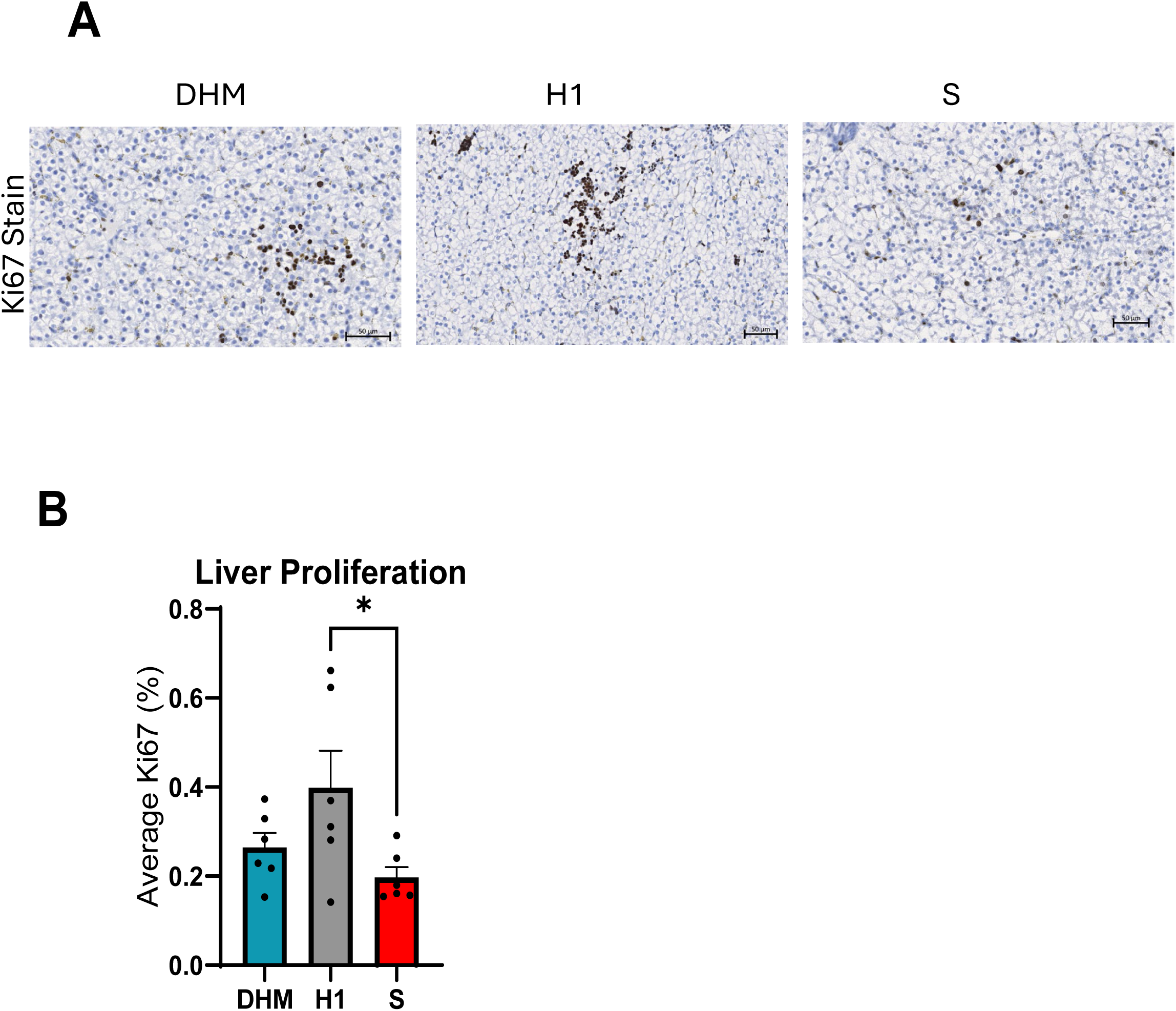
Liver histology Ki67 staining in piglets on d 10. A) Liver cell proliferation as measured by representative images of Ki67 staining from piglets fed either donor human milk (DHM), novel infant formula (H1; Boston, MA, US), or term infant formula (S; Similac; Abbott, Chicago, IL) on d10; Scale bar, 50 µm. B) Quantification of mean percentage Ki67 staining in liver tissue in each treatment group. *, *P* < 0.05

### Plasma, Distal Ileum, and Liver Cytokines

Several cytokines that were tested were below the level of detection for all treatments and therefore not included in the results. There were no differences (*P* > 0.05) in the plasma (**Figure 6A–F**) and liver (**Figure 7A-I**) across all treatments for the detectable cytokines. However, there were some differences across treatments in the distal ileum (**Figure 8A-K**). IL-4 (**Figure 8F**) was elevated in piglets that received S compared to DHM. Additionally, in the distal ileum, H1 had more IL-8 than both DHM and S-fed piglets (**Figure 8H**). Overall, these were the only two differences in all the cytokines that were measured.

**Figure 6.**
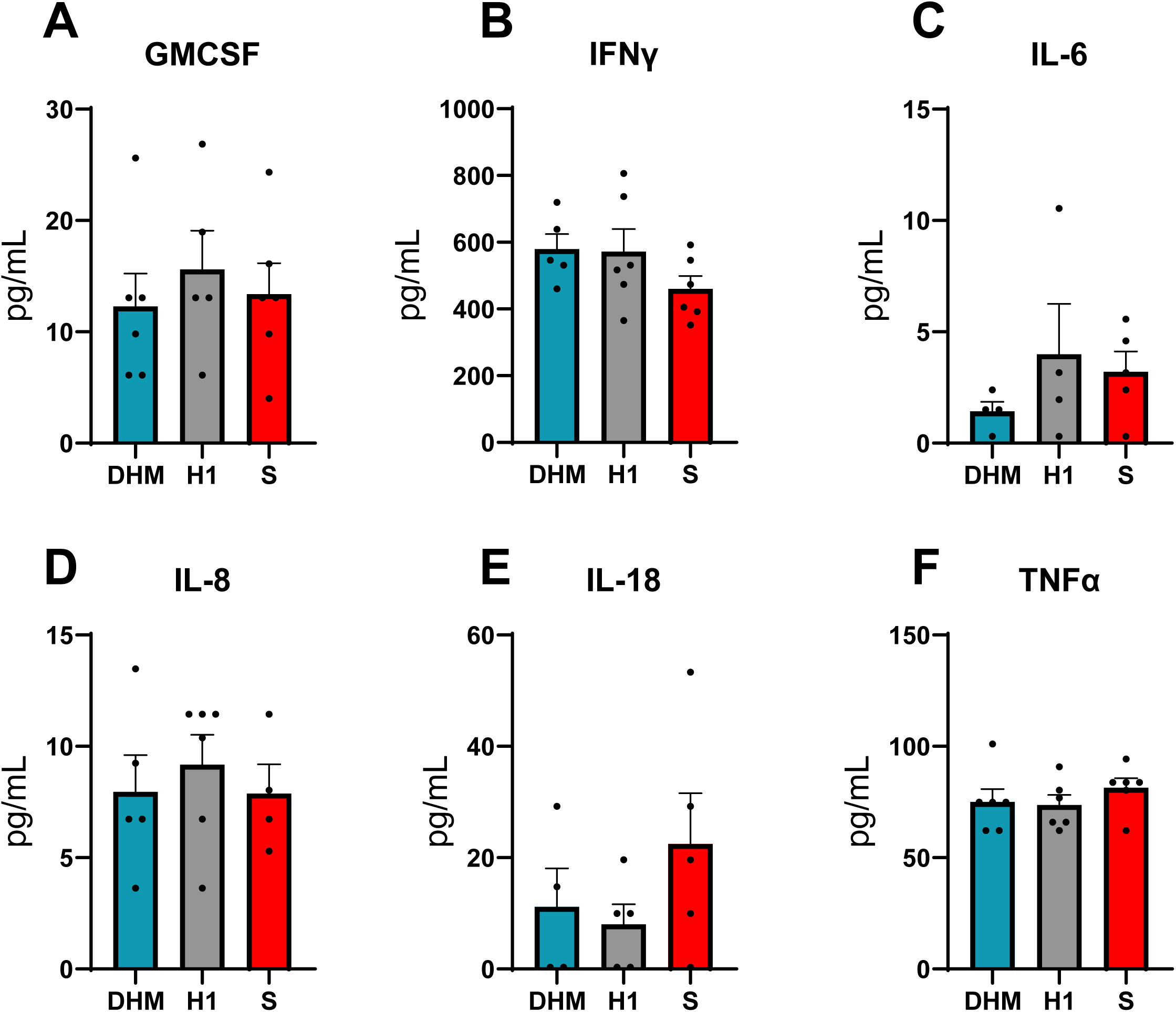
Plasma cytokines of piglets on d 10. A-K) individual analysis of cytokines found in circulation on d 10 of piglets fed donor human milk (DHM), novel infant formula (H1; Boston, MA, US), or term infant formula (S; Similac; Abbott, Chicago, IL); GMCSF, granulocyte-macrophage colony-stimulating factor; IFNγ, interferon gamma; IL, interleukin; TNFα, tumor necrosis factor-alpha.

**Figure 7.**
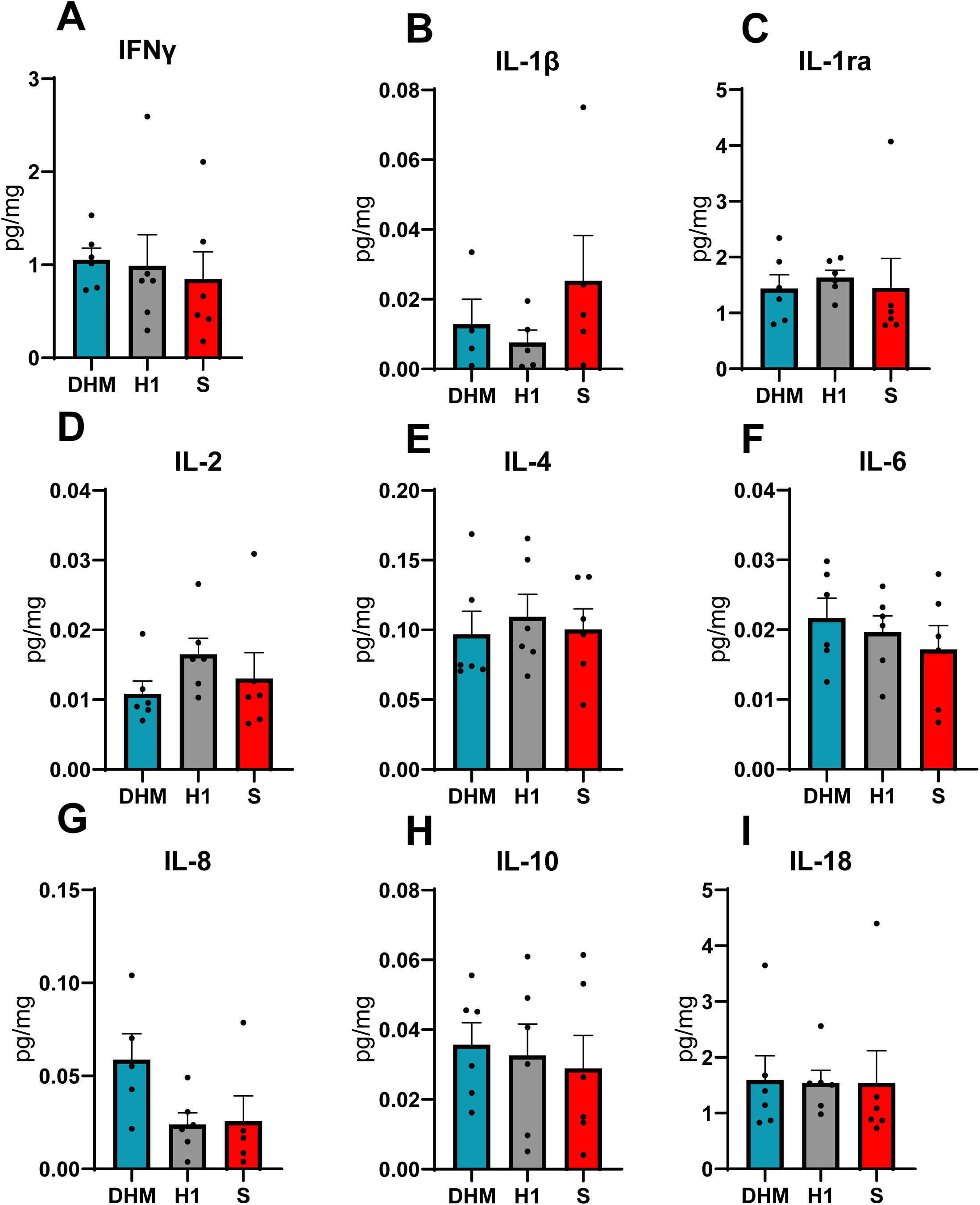
Liver cytokines of piglets on d 10. A-K) individual analysis of cytokines found in the liver on d 10 of piglets fed donor human milk (DHM), novel infant formula (H1; Boston, MA, US), or term infant formula (S; Similac; Abbott, Chicago, IL); IFNγ, interferon gamma; IL, interleukin; IL-1ra, Interleukin-1 receptor antagonist.

**Figure 8.**
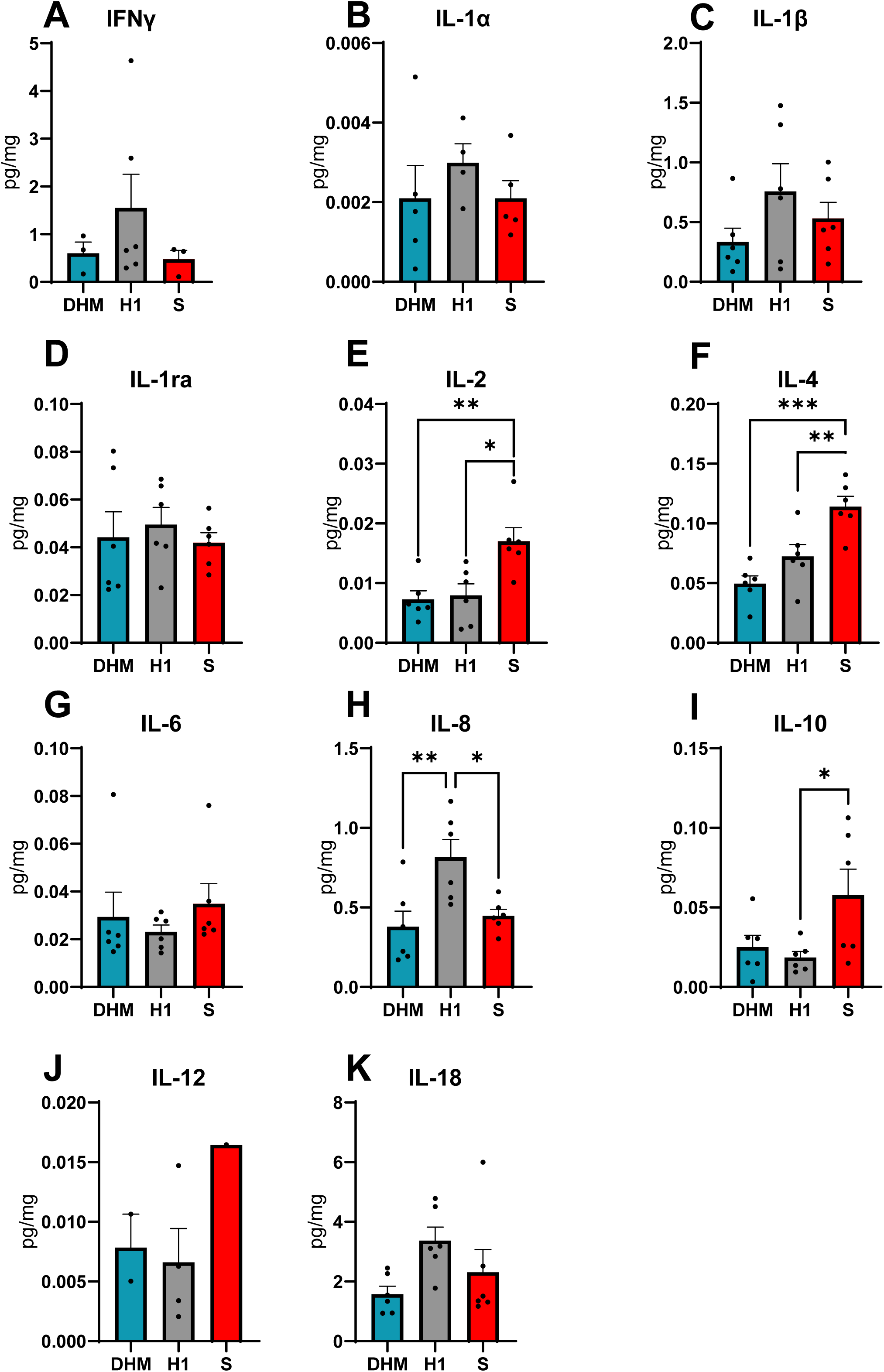
Distal ileum cytokines of piglets on d 10. A-K) individual analysis of cytokines found in the distal ileum on d 10 of piglets fed donor human milk (DHM), novel infant formula (H1; Boston, MA, US), or term infant formula (S; Similac; Abbott, Chicago, IL); IFNγ, interferon gamma; IL, interleukin; IL-1ra, Interleukin-1 receptor antagonist. *, *P* < 0.05; **, *P* < 0.01; ***, *P* < 0.001

### Plasma Amino Acid Concentrations

Plasma amino acid concentrations at day 10 showed select differences among treatment groups (**Figure 9A and 9B**). The essential amino acid histidine was lower (*P* < 0.05) in piglets that received S, compared to piglets that received DHM and H1, however; methionine is higher (*P* < 0.05) in piglets that received S than piglets that received DHM or H1. Phenylalanine in H-fed piglets was higher (*P* < 0.05) than DHM-fed piglets. H1 fed piglets also had higher (*P* < 0.05) Lysine than DHM piglets, and S fed piglets had higher (*P* < 0.05) Lysine than DHM piglets. Among nonessential amino acids (**Figure 9B**), H1 fed piglets had higher (*P* < 0.05) arginine and tyrosine than DHM or S. Piglets fed H1 also had higher (*P* < 0.05) glutamine than piglets fed S. Lastly, H1 piglets had higher (*P* < 0.05) serine and ornithine than DHM piglets.

**Figure 9.**
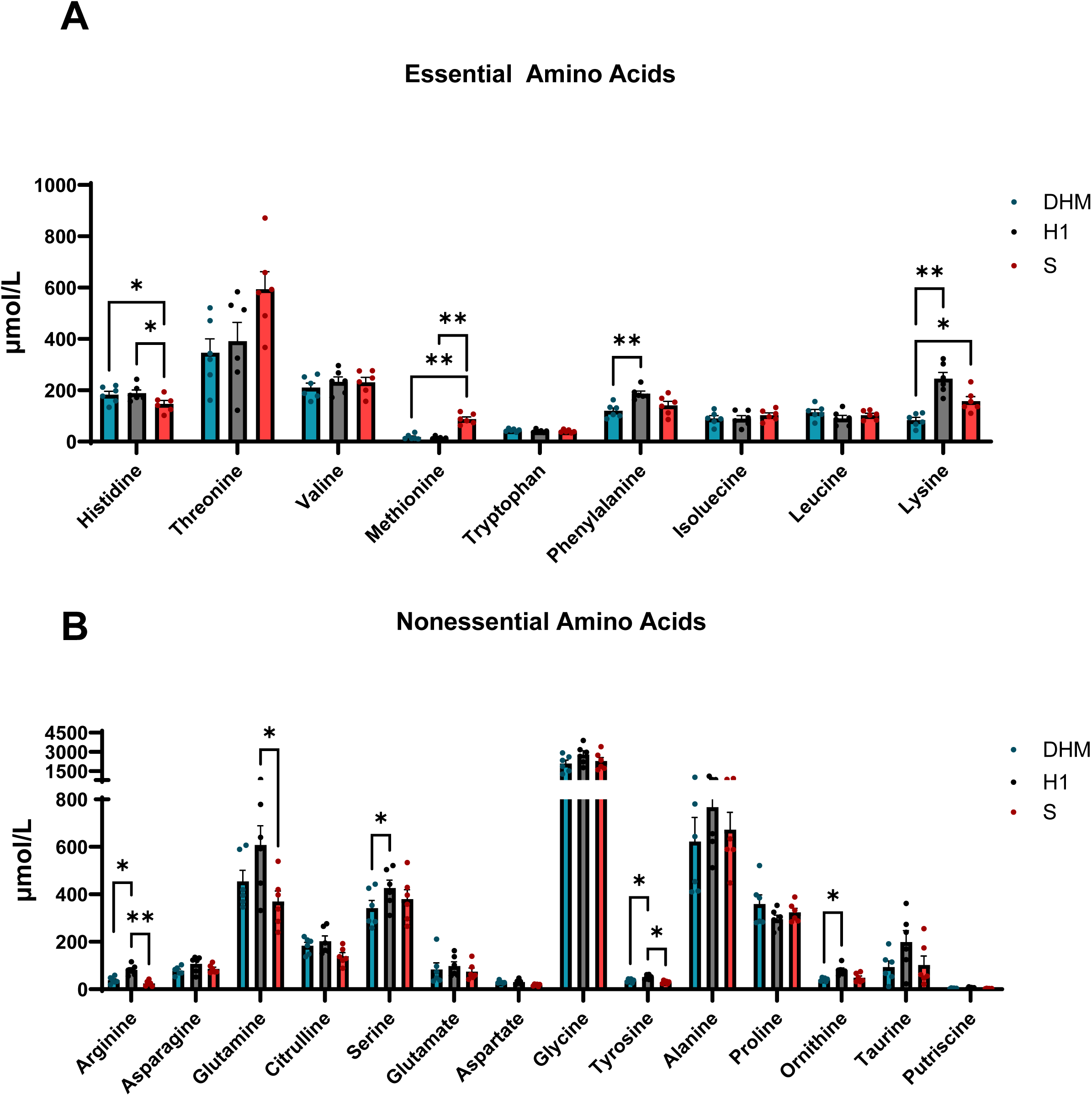
Circulating amino acid concentrations of piglets on d 10. A) Essential amino acid values in the serum of piglets fed donor human milk (DHM), novel infant formula (H1; Boston, MA, US), or term infant formula (S; Similac; Abbott, Chicago, IL) on d 10 B)) Nonessential amino acid values in the serum of pigs fed DHM, H1, or S on d 10. *, *P* < 0.05; **, *P* < 0.01

### Esterase Activity

The esterase activity (**Figure 10A**) of serum albumin was measured to confirm the proper folding of the protein and compare it to the recombinant and native protein activity. The recombinant protein enzyme activity is roughly 47% lower than the native protein. The recombinant protein has a higher Km (∼290%) and a similar V_max_ compared to the native protein.

**Figure 10.**
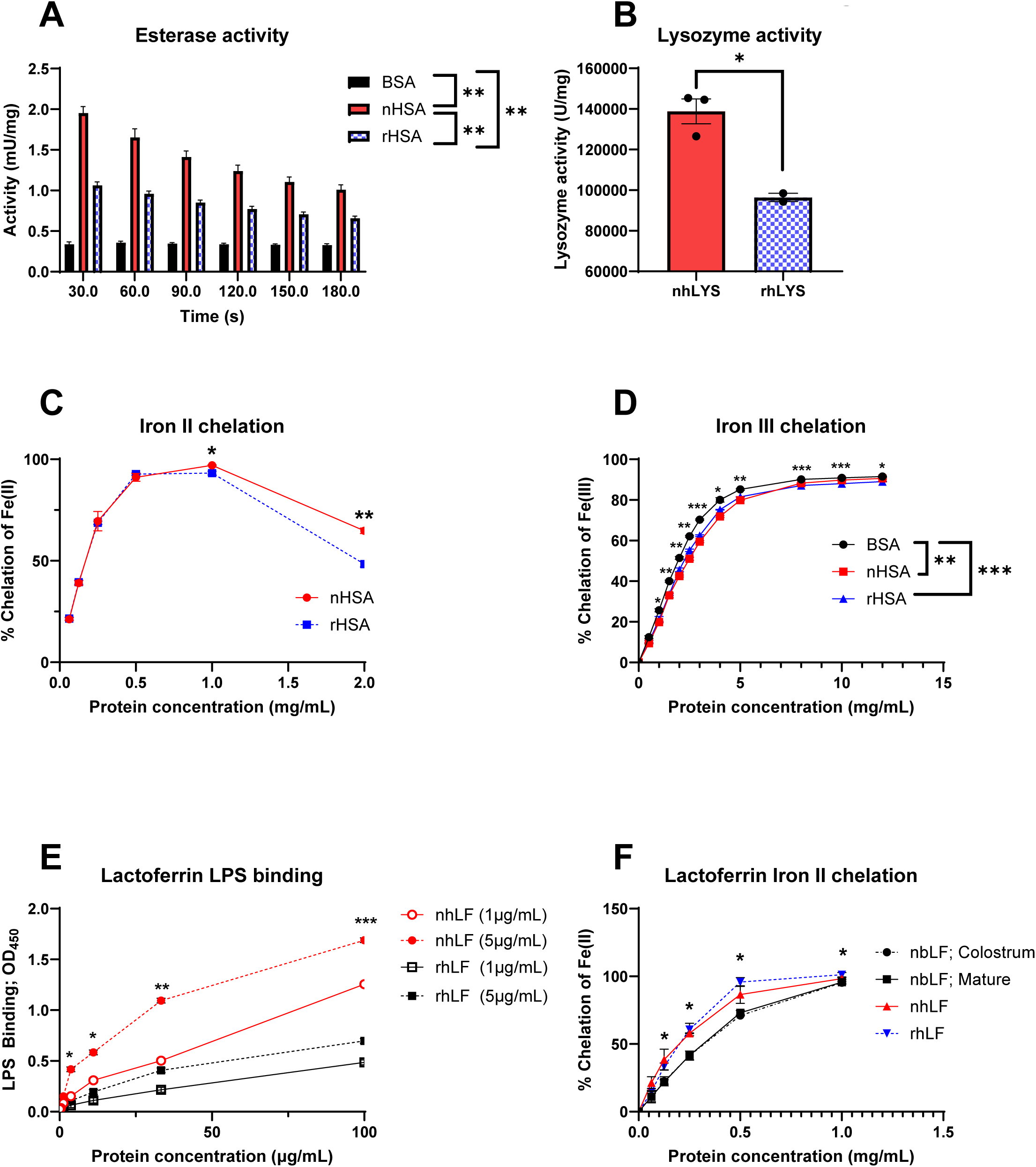
Recombinant human protein analysis compared to native human proteins and bovine proteins. A) Esterase activity was significantly different at all time points, with native human serum albumin (nHSA) being significantly higher than bovine serum albumin (BSA) and recombinant human serum albumin (rHSA). B) Lysozyme activity of native human lysozyme (nhLYS) and recombinant human lysozyme (rhLYS). C) Human serum albumin chelation of iron II was similar until 1.0 mg/mL and 2.0 mg/mL of protein, at which point nHSA was significantly higher than rHSa. D) BSA, nHSA, and rHSA chelation of iron III. BSA had a higher percent of chelation of iron III compared to both nHSA and rHSA. E) Lactoferrin Lipopolysaccharide (LPS) binding ability for either native human lactoferrin (nhLF) or recombinant human lactoferrin (rhLF). Starting at 3.7 µg/mL of protein, nhLF at 5 µg/mL is binds more LPS than other proteins, and this remains true through higher protein concentrations. At 100 µg/mL protein concentration, nhLF at 1 µg/mL binds more LPS than both rhLF. There are never any differences between 1 µg/mL and 5 µg/mL of rhLF. F) Lactoferrin iron II chelation assessment of native bovine lactorferrin (nbLF), nhLF, and rhLF. Both nhLF and rhLF have a higher percent iron II chelation than nbLF from colostrum and mature milk. *, *P* < 0.05; **, *P* < 0.01; ***, *P* < 0.001

Throughout the kinetic assay, there is an effect of time (*P* < 0.0001) in which activity decreased from the start of the assay to the end. The highest activity points were: 60s for bovine serum albumin (BSA); 30s for native human serum albumin (nHSA); and 30s recombinant human serum albumin (rHSA). At every time point, nHSA has the highest activity (*P* < 0.05) compared to BSA and rHSA, and BSA activity is lower (*P* < 0.05) than nHSA and rHSA activity.

### Lysozyme Activity on Isolated Protein

Lysozyme activity was measured by breaking down the peptidoglycan wall of bacteria (**Figure 10B**). The lysozyme activity of recombinant human lysozyme is lower (*P* < 0.05), about 31% lower, than native human lysozyme (138,817 U/mg vs 96,392 U/mg, native human lysozyme activity vs recombinant human lysozyme activity, respectively). This shows that the recombinant human lysozyme activity is still present, but at a lower rate than the native protein.

### Iron II and III Binding on Isolated Serum Albumin

The ability to chelate iron II and carry it across the gut epithelial barrier is important to ensure that infants are receiving the proper micronutrients. The native and recombinant HSA ability to chelate iron II was similar up to 1 mg/mL (**Figure 10C**). Iron II had the highest (*P* < 0.05) chelation percentage at 1 mg/mL protein concentration for nHSA, while rHSA had the highest (*P* < 0.05) percent chelation at 0.5 mg/mL and 1 mg/mL protein concentration. Higher concentrations of the proteins result in the recombinant protein having a lower percentage of chelation of iron II. However, there were minimal differences in the percentage of chelation of iron III between BSA, nHSA, and rHSA (**Figure 10D**).

### Lactoferrin LPS Binding

Measuring the level of interaction between the recombinant and native human lactoferrin and the LPS on Escherichia coli can help to confirm if the antimicrobial activity from the recombinant protein is similar to the native protein. There are no differences in the LPS binding capability of native human lactoferrin (nhLF) and recombinant human lactoferrin (rhLF) when looking at LPS concentrations of 0.00 through 1.23 µg/mL (**Figure 10E**). When the LPS concentration is increased to 3.7 µg/mL, nhLF (5 µg/mL) has the highest LPS binding, indicated by a higher OD_450_ (*P* < 0.05) compared to nhLF (1 µg/mL), rhLF (µg/mL), and rhLF (µg/mL). The same is true at 33.3 µg/mL with hnLF (1 µg/mL) also having a higher LPS binding than rhLF (1µg/mL). Finally, all this holds true at 100 µg/mL LPS with also nhLF (1 µg/mL) having a higher LPS binding than rhLF (5 µg/mL). Overall, recombinant lactoferrin has around 60% less binding capacity than the native protein.

### Lactoferrin Iron II

There are two iron binding sites on lactoferrin, which have a higher affinity for iron III, but it is assumed that iron II can also bind to these sites (22). Thus, the amount of iron II chelated by lactoferrin was indirectly measured and compared to the native and recombinant protein activities. There were no differences (*P* > 0.05) in the percent chelation of iron II when there were 0 and 0.063 mg/mL of the lactoferrin from any source. Starting at 0.125 mg/mL of lactoferrin, there is a difference (*P* < 0.05) in the percent chelation of iron II for native bovine colostrum lactoferrin (nbLF; colostrum) and the recombinant human lactoferrin (rhLF), with rhLF having a higher percent chelation. rhLF also had a higher percent chelation (*P* < 0.05) compared to the native bovine mature milk lactoferrin (nbLF; mature). The same difference (*P* < 0.05) held true at 0.250 mg/mL of lactoferrin protein. Additionally, the native human lactoferrin (nhLF) also had a higher (*P* < 0.05) percent chelation of iron II compared to nbLF; colostrum and nbLF; mature. At 0.500 mg/mL lactoferrin, the results were the same as at 0.125 mg/mL, rhLF had a higher (*P* < 0.05) percent chelation of iron II than nbLF; colostrum and nbLF; mature. At 1.000 mg/mL lactoferrin, the percent chelation of iron II for rhLF was greater (*P* < 0.05) than nbLF; colostrum and nhLF had a greater (*P* < 0.05) chelation of iron II than nbLF; mature. Overall, the native and recombinant human lactoferrin were similar to native bovine colostrum and mature milk lactoferrin.

## Discussion

The present study compared a novel proof-of-concept infant formula (H1) containing recombinant human proteins (albumin, lactoferrin, and lysozyme) against donor human milk (DHM) and a standard commercial term infant formula (S) utilizing a translational term neonatal piglet model. Bridging the nutritional and functional gap between human breast milk and bovine-based infant formulas remains a significant challenge in pediatric nutrition. The primary finding of this study is that the H1 formula was not noninferior to DHM and S in supporting overall somatic growth and maintaining small intestinal integrity over a 10-day neonatal piglet feeding period. Moreover, we found no differences in small intestinal or systemic cytokines concentrations in piglets and our in vitro evidence showed that the recombinant human proteins used retained their functions compared to their respective native human and bovine proteins.

### Growth and Organ Development

There were no significant differences in final body weight or body weight gain among the dietary treatments. This indicates that the macronutrient profile of H1, which provides similar levels of calories, carbohydrates, protein, and fat to standard term infant formula, is sufficient to support normal neonatal growth trajectories comparable to human milk. It is important to note that the protein intake fed to piglets in this study was well below that required (∼15 g protein/kg per day) to achieve more normal growth (50-75 g/kg per day) in sow-reared and artificially-reared formula- and donor human milk fed piglets fed to meet the pig nutrient requirements (13, 23). We chose this approach to approximate slower daily growth rates in human term (5-10 g/kg) and preterm infants (10-15 g/kg) while using the piglets higher growth potential to detect difference between the test diets. This was reflected in a daily growth rates among the three treatment groups ranging from 20-30 g/kg for term neonatal pigs in the first 10 days postnatal. Notably, piglets fed the H1 formula exhibited significantly smaller relative liver and stomach weights compared to those receiving DHM or S. The reduced relative stomach weight may be attributed to the physical characteristics of the H1 formula; its more aqueous nature may facilitate easier gastric emptying and reduce compliance limiting its physical expansion. Similarly, the smaller relative liver weight suggests that the specific nutrient and free amino acid matrix of H1 exerts a lower metabolic workload on the liver. Despite these treatment difference the relative weights of liver and intestine were within the normal range for sow-fed or artificially reared piglets (23).

### Hematology, Systemic Immunity, and Intestinal Histology

Evaluation of systemic immunity via complete blood counts showed no significant differences in white blood cells (WBC), neutrophils, lymphocytes, or eosinophils across the DHM, H1, and S groups, indicating a comparable systemic immune response to the diets. However, compared to previously established reference intervals for 5-day-old piglets (21), the S-fed group showed elevated WBC and neutrophil counts. While this hints at a potential mild systemic immune response in S-fed pigs, all piglets in this study received iron, whereas those in the reference study did not receive an iron injection, which is critical to piglet growth in the neonatal period.

At the intestinal mucosal level, mucin content in the distal ileum was comparable across all treatments. This suggests that the formulas did not negatively alter goblet cell mucin production or fundamental absorptive barrier functions, and that nutrients are absorbed uniformly across diets without triggering a compensatory mucin response. Interestingly, liver cell proliferation, marked by Ki67 staining, was significantly higher in H1-fed piglets compared to S-fed piglets. This localized cellular activity presents an intriguing contrast to the lower relative liver weights observed in the H1 cohort and warrants further investigation to define its possible physiological benefit.

### Cytokine Profiling and Localized Inflammation

Histological inflammation scores were generally low and similar across all treatments, confirming that H1 does not induce excessive intestinal inflammation in a healthy neonatal model. Cytokine profiling further confirmed an absence of systemic or hepatic inflammation, as plasma and liver cytokine levels were either undetectable or showed no significant differences. In the distal ileum, however, notable localized differences emerged. Piglets fed standard formula (S) exhibited elevated IL-4 compared to the DHM group. Because IL-4 is integral to macrophage regulation, T-cell development, and allergic responses, its elevation may reflect a mild hypersensitivity or allergic inflammatory reaction to bovine proteins in the standard formula (24). Conversely, H1-fed piglets exhibited elevated IL-8 and IL-18 in the distal ileum compared to other groups. While IL-18 possesses proinflammatory properties, its isolated elevation—without corresponding systemic markers—may indicate early localized immune priming or a transient response to environmental microbes rather than pathogenic inflammation (25).

### Amino Acid Bioavailability

Optimal neonatal growth and development relies on amino acid bioavailability of dietary protein. The H1 formula demonstrated excellent protein digestibility and systemic amino acid delivery. Plasma analysis at day 10 revealed that H1-fed piglets maintained equivalent or significantly higher concentrations of several amino acids compared to both the DHM and S groups. Specifically, H1-fed piglets showed elevated circulating levels of essential amino acids like phenylalanine and lysine, as well as nonessential amino acids including arginine, tyrosine, glutamine, serine, and ornithine. The S formula yielded lower levels of histidine, but higher methionine compared to DHM and H1. Overall, these data confirm that the free amino acids and recombinant proteins in H1 are highly bioavailable, effectively supporting circulating amino acid pools without acting as a limiting factor for protein deposition or metabolic function.

### Recombinant Protein Bioactivity

A major innovation of the H1 formula is the inclusion of functional recombinant human proteins to mimic the diverse bioactive and immunological components of human milk. *In vitro* functional assays demonstrated that while the enzymatic and binding activities of these recombinant proteins were sometimes marginally lower than their native human counterparts, they matched or outperformed standard commercial bovine proteins. Serum albumin has a role not only in blood osmolarity, but can also be a transport protein for fatty acids and has binding sites for several metabolites, such as medium-chain fatty acids in humans (26). Additionally, serum albumin has catalytic activity for organic molecules. It has been shown to have some esterase activity, as human serum albumin has been documented to hydrolyze *p*-nitrophenol on esters (27). The esterase activity of recombinant human serum albumin (rHSA) was approximately 47% lower than native human serum albumin (nHSA) but remained significantly higher than bovine serum albumin (BSA), indicating its retained capacity to assist in organic molecule hydrolysis, protein folding, and toxic substance binding. Furthermore, rHSA showed equivalent iron II chelation to nHSA at lower concentrations and equivalent iron III chelation across all tested concentrations. This chelation is critical for infant immunity, as it restricts bacterial iron utilization and prevents the generation of reactive hydroxyl radicals.

Serum albumin and lactoferrin can chelate iron which is critical for the immune system as it prevents the formation of hydroxyl radicals and prevents bacteria from using iron for growth (28). Without the sequestration of iron, bacteria use it for growth and virulence. Iron III is poorly water-soluble and binds to proteins, such as transferrin, to be stable and have bioavailability (29). Meanwhile, iron II is water-soluble and has high reactivity, leading the generation of hydroxyl radicals (29). The assessment of lactoferrin activity is important because it constitutes 15-20% of the total protein content in human breast milk (5). Lactoferrin is antibacterial, antiviral, and anti-inflammatory (30). Recombinant human lactoferrin (rhLF) bound lipopolysaccharide (LPS) at approximately 60% the capacity of native human lactoferrin, confirming its functional antimicrobial properties. Importantly, rhLF demonstrated superior iron II chelation compared to native bovine lactoferrin sourced from both colostrum and mature milk. Similarly, recombinant human lysozyme (rhLYS) maintained structural degradation capabilities against bacterial peptidoglycan, albeit at a ∼31% lower activity rate than native human lysozyme. This highlights the physiological and functional advantage of utilizing human-identical recombinant proteins over bovine alternatives in infant nutrition.

A primary strength of this study is the integration of multiple functional recombinant proteins (rhHSA, rhLYS, rhLF) into a single formula matrix, as opposed to prior studies which heavily relied on evaluating a single transgenic protein. Secondly, this study used a translational neonatal piglet model paired with comprehensive histological, immunological, and functional protein assays. Prior animal models have shown that rhLF and rhLYS inhibit gut microbes, specifically *Escherichia coli* and promoting beneficial *Bifidobacterium* and *Lactobacillus* species, however this study did not evaluate the impact of H1 on microbial populations.

### Conclusion

The current study showed that a novel infant formula, fortified with human recombinant proteins, is noninferior to donor human milk and standard infant formula in supporting short-term neonatal growth and intestinal histology. The enhanced bioavailability of amino acids, coupled with the retained bioactivity of recombinant human lactoferrin, albumin, and lysozyme, suggests that this “humanized protein” formula represents a viable, hypoallergenic, next-generation alternative to bovine-based infant formulas. Future research should investigate its long-term metabolic effects and its precise role in shaping the infant gut microbiota.

## Acknowledgements

The authors would like to acknowledge Inka Didelija, Liwei Cui, and Xiaoyan Chang for their hard work in completing laboratory assays and assisting with piglet care, which contributed significantly to the completion of this project. We also acknowledge Robert DiGregorio and Vanessa Castagna for their consultation on the development of the novel formula composition used in the study.

## The author’s responsibilities were as follows

SE, VHM, BS, DB - Designed the study; SE, VHM, GH, JL, MWSdS, MR, CV, GG, BS - Conducted research; GH, JL, JR, MWSdS, MR - Provided essential reagents and essential materials; SE - Analyzed data; SE, GH, JL, MWSdS, MRS - Wrote paper; VHM, GH, JL, JR, MWSdS, MRS, CV, GG, BS, DB - Edited manuscript; DB - Primary responsibility for final content.

## Conflicts of Interest

Authors GH, JL, JR, MWSdS, MRS and DA work at Harmony Baby Nutrition, which funded a large part of the research presented herein.

## Declaration of Generative AI and AI-assisted technologies in the scientific writing process

AI was not used in the production of this manuscript.

## Data Sharing

Data described in the manuscript, code book, and analytic code will be made available upon request, pending approval from authors.

## Disclaimer

Harmony Baby Nutrition funded a large part of the work presented in this manuscript.

## Sources of support

This work was supported in part by federal funds from the USDA, Agricultural Research Service under Cooperative Agreement Number 58-6250-6-001, and the National Institutes of Health Grant DK-094616 and Texas Medical Center Digestive Diseases Center (NIH Grant P30 DK-56338)(D.B). The work was supported by a grant from Harmony Baby Nutrition. S Elefson was supported by training fellowship grants from the National Institutes of Health Grant T32-DK07664. V Melendez Hebib was supported by institutional training grant T32 GM136554. C. Vonderohe was supported by a training fellowship and grants from the National Institutes of Health Grant T32-DK07664 and K08 DK135845. Guthrie was supported by a training grant from the National Institutes of Health Grant K01 DK129408

## Abbreviations

BSA: Bovine Serum Albumin
CXCL8: C-X-C Motif Chemokine Ligand 8
CBC: Complete Blood Count
DHM: Donor Human Milk
ELISA: Enzyme-Linked Immunosorbent Assay
GM-CSF: Granulocyte-Macrophage Colony-Stimulating Factor
HLF: Human Lactoferrin
HMOs: Human Milk Oligosaccharides
HAS: Human Serum Albumin
IFN-γ: Interferon-Gamma
IL-1α: Interleukin-1 Alpha
IL-1β: Interleukin-1 Beta
IL-1ra: Interleukin-1 Receptor Antagonist
IL-2: Interleukin-2
IL-4: Interleukin-4
IL-6: Interleukin-6
IL-8: Interleukin-8
IL-10: Interleukin-10
IL-12/IL-23(p40): Interleukin-12/Interleukin-23 Subunit p40
IL-18: Interleukin-18
LPS: Lipopolysaccharide
nbLF: Native Bovine Lactoferrin
nhLF: Native Human Lactoferrin
nhLYS: Native Human Lysozyme
nHSA: Native Human Serum Albumin
H1: Novel Investigational Formula
PBS: Phosphate-Buffered Saline
PBST: Phosphate-Buffered Saline with Tween 20
rh: Recombinant Human
rhLF: Recombinant Human Lactoferrin
rhLYS: Recombinant Human Lysozyme
rHSA: Recombinant Human Serum Albumin
S: Term Infant Formula
TBS: Tris-Buffered Saline
TNF-α: Tumor Necrosis Factor-Alpha
WBC: White Blood Cells

## Notes

### Competing Interest Statement

The authors have declared no competing interest.

